# Gene family expansions and nodule-specific expression patterns reveal the recruitment of Beta-Glucosidases and Cytochrome P450 genes to nodulation in soybean

**DOI:** 10.1101/2025.07.03.662952

**Authors:** Cláudio Benício Cardoso-Silva, Gabriel Quintanilha-Peixoto, Dayana Kelly Turquetti-Moraes, Fabrício Almeida-Silva, Thiago M. Venancio

## Abstract

Nodulation is a symbiotic interaction between legumes and rhizobia that is essential for biological nitrogen fixation and regulated by complex gene networks, which are partially shaped by whole-genome duplication (WGD) events. Although many core nodulation genes have been identified, the contribution of lineage-specific gene expansions and WGD-derived duplicates remains poorly understood. In this study, we performed a genome-wide analysis of gene family expansions in legumes and integrated expression data from over 5,000 soybean RNA-Seq samples to identify genes involved in nodulation. Our analysis not only recovered well-characterized nodulation genes among the most expanded families but also revealed several differentially expressed genes in nodules, including nodulins, aspartic proteases, NLP proteins, and GRAS transcription factors. Notably, we identified previously unreported β-glucosidase and cytochrome P450 genes with high nodule-specific expression, suggesting their involvement as novel components of the nodulation machinery.

## INTRODUCTION

Nodulation is a complex and highly regulated process in which beneficial bacteria, collectively referred to as rhizobia, establish symbiotic relationships with legume roots, forming specialized structures known as nodules. These nodules house rhizobia and enable the fixation of atmospheric nitrogen into forms usable by the plant (Goto et al., 2022). The establishment and maintenance of this symbiosis are governed by intricate molecular signals and gene networks that regulate interactions between legumes and rhizobia (Roy et al., 2020). Legume nodules are typically classified as determinate or indeterminate based on the activity of the apical meristem. Determinate nodules, as found in soybean, common bean, and cowpea, lose meristematic activity shortly after initiation. In contrast, indeterminate nodules, characteristic of species such as pea, alfalfa, and clover, retain an active apical meristem throughout development (J. Liu & Bisseling, 2020). Although nodulation is widespread among legumes, certain genera, including *Cercis* (Granqvist et al., 2015)*, Nissolia* (Griesmann et al., 2018), and *Senna* (Arenque et al., 2014), have lost nodulation capacity.

Whole-genome duplication (WGD) has played a substantial role in shaping the gene complement associated with nodulation (Li et al., 2013). WGDs have occurred repeatedly throughout angiosperm evolution (Renny-Byfield & Wendel, 2014; D. E. Soltis et al., 2009; P. S. Soltis et al., 2015), acting as a major force for evolutionary innovation, particularly in plants (Heslop-Harrison et al., 2023; Hirakawa, 2022; P. S. Soltis & Soltis, 2016; Tao et al., 2023; Tronconi et al., 2020; Van de Peer et al., 2009, 2017). It is estimated that approximately 65% of the plant genes have arisen through WGD events (Panchy et al., 2016). In addition to its relevance for nodulation, the retention of WGD-derived gene duplicates contributes to the genomic plasticity of soybean, enhancing its adaptability and resilience to environmental challenges (Ebadi et al., 2023). Although many duplicate genes are eventually lost over time, some are retained and undergo subfunctionalization or neofunctionalization (Sandve et al., 2018; Tautz & Domazet-Lošo, 2011), driving the expansion and diversification of gene families at both sequence and expression levels (Landis et al., 2018; Nei & Rooney, 2005). Soybean, in particular, has experienced two rounds of WGD in its evolutionary history, resulting in extensive gene redundancy and diversification (Almeida-Silva & Van de Peer, 2025; Schmutz et al., 2010).

These duplication events have provided the raw material for the emergence and functional diversification and refinement of nodulation-related genes (De Mita et al., 2014; T. Liu et al., 2024; Op den Camp et al., 2011; Powell & Doyle, 2015; Rutten et al., 2020, 2020). Key components of the nodulation machinery include nodule inception (NIN) (J. Liu & Bisseling, 2020), nodulation-specific lipochitooligosaccharide (LCO) receptors (Geurts & Huisman, 2024), IAA carboxyl methyltransferase 1 (Goto et al., 2022), a legume-specific CLAVATA/ESR-related factor (Mens et al., 2021), PLAT domain proteins (Trujillo et al., 2019), nodule-specific cysteine-rich peptides (Mergaert et al., 2003), and calmodulin-like proteins (J. Liu et al., 2006). These gene families collectively orchestrate the molecular signaling and developmental pathways required for effective nodulation. Scientific advances in understanding nodulation have significantly contributed to improved soybean productivity and reduced dependence on synthetic nitrogen fertilizers (Xu et al., 2020). Recent studies have uncovered novel transcription factors (TFs) and elucidated signaling networks central to this symbiosis (Amin et al., 2023; Chen et al., 2020; L. Wang et al., 2019; Yun et al., 2022). Nonetheless, important gaps persist, especially regarding the recruitment of lineage-specific genes and the evolutionary impact of WGD and gene family expansions on nodulation.

To address such knowledge gaps, we conducted a comprehensive genome-wide analysis of gene family expansions across legumes, integrating these data with expression profiles from over 5,000 soybean RNA-Seq samples. Our primary goal was to identify novel genes potentially involved in nodulation, with a focus on targets that could support crop improvement and sustainable agriculture. Our findings confirmed previous reports by highlighting the involvement of key genes associated with nodulation among the top 100 expanded gene families. Moreover, we identified several genes with differential expression in nodules, including nodulins, aspartic proteases, NLP proteins, and GRAS TFs. Notably, we also discovered previously unreported β-glucosidase and Cytochrome P450 genes that are highly expressed in nodules, suggesting new components of the nodulation machinery in soybean.

## MATERIALS AND METHODS

### Inference of orthologous families, expansions, and contractions

To estimate gene family expansion and contraction rates in legumes, we selected the predicted protein sequences of 35 species across three orders within Fabidae (Fabales, Fagales, and Rosales) (Table S1), including four non-nodulating species (*Cercis canadensis*, *Nissolia schottii*, *Senna tora*, and *Juglans regia*). Redundant proteins were removed using CD-HIT v4.8.1 (Fu et al., 2012) with a coverage threshold of 99%. Orthogroup (OG) inference was performed using OrthoFinder v2.5.1 (Emms & Kelly, 2019), with Diamond v0.9.24 (Buchfink et al., 2015) for BLASTp (e-value ≤ 1e-5) and MAFFT v7.505 (Katoh et al., 2019) for multiple sequence alignment. A phylogenomic tree was reconstructed using single-copy orthologs identified by OrthoFinder, with IQ-TREE v2.1.2 (Minh et al., 2020) employed for maximum likelihood tree inference, incorporating automated model selection and 1000 bootstrap replicates. Tree visualization was carried out using the R package *ggtree* v3.4.0 (Yu et al., 2017). Single-copy orthologs were selected using STAG (Emms & Kelly, 2019), and tree rooting was performed using STRIDE (Emms & Kelly, 2017). Gene family expansions and contractions were inferred using CAFE5 v5.0 (Mendes et al., 2021), which applies a likelihood-based stochastic birth-death model on gene count data and a time-calibrated species phylogeny. To reduce bias from extreme OG size variation, CAFE built-in preprocessing step was used to filter OGs with high variance in gene counts across species. The ultrametric phylogenetic tree required by CAFE was generated using *in-house* scripts, incorporating fossil calibration points obtained from the TimeTree database (Kumar et al., 2017). Tandem and segmental gene duplications were identified using MCScanX (Y. Wang et al., 2012), enabling classification of the mechanisms underlying gene family expansion.

### Gene expression analysis

We performed gene expression analyses using data from the Soybean Expression Atlas (Almeida-Silva & Venancio, 2023; Machado et al., 2020), which comprises over 5,000 RNA-Seq samples spanning a wide range of tissues (available at https://soyatlas.venanciogroup.uenf.br). For comparative purposes, we also analyzed RNA-Seq data for *Medicago truncatula* retrieved from NCBI BioProjects PRJEB43588 and PRJNA832760, each comprising 24 samples. Quality control and adapter trimming of the *M. truncatula* RNA-seq reads was carried out using Trimmomatic v0.36 (Bolger et al., 2014). Transcript abundance across nine tissues (coat, embryo, epidermis, inflorescence, leaf, nodule, root, seed, and stem) was estimated using Kallisto v0.50.0 (Bray et al., 2016) and normalized in transcripts per million (TPM). Due to the high expression variance across samples from different experiments, we computed the geometric mean of TPM values to represent average gene expression in each tissue (Vandesompele et al., 2002).

### Functional annotation of candidate genes

One selected gene family (ORTHO05D000033) was functionally annotated using InterPro (Jones et al., 2014), applying an e-value cut-off of 1e-10. To group functionally similar members into isofunctional clusters, defined as sets of genes with similar predicted functions, we constructed a sequence similarity network (SSN) (Gerlt et al., 2015). The SSN was further enhanced and visually explored using Cytoscape v3.9.1 (Shannon et al., 2003).

### Gene duplication classification

Gene duplication types in the soybean genome were identified using the Doubletrouble database (Almeida-Silva & Van de Peer, 2025). For each duplicated gene pair, we computed the Synonymous (Ks) and non-synonymous (Ka) substitution rates using the KaKs_Calculator3.0 (Zhang, 2022), applying the maximum-likelihood Goldman-Yang method. The duplication time (T) for each gene pair was estimated using the formula:

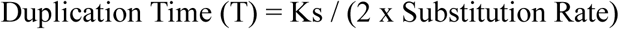

A substitution rate of 6.5 × 10^−9^ substitutions per synonymous site per year was used, as previously reported (Gaut et al., 1996; Tang et al., 2010).

## RESULTS AND DISCUSSION

### Gene family expansions and contractions in legumes are consistent with WGD events

Our findings indicate that gene family expansions within the genus *Glycine* are closely associated with WGDs in their evolutionary history. We analyzed 1,365,329 protein-coding genes from 35 legume species, identifying 49,785 orthologous gene families. The number of non-redundant genes per species, defined by collapsing sequences sharing ≥99% similarity (see Methods for details), ranged from 20,946 in *Lablab purpureus* to 67,150 in *Arachis hypogaea*. *Glycine max* and *Glycine soja* contained 56,044 and 44,969 non-redundant genes, respectively (Figure 1, Figure S1). Additional information on the selected species is available in Table S1. On average, 94.7% of the genes were assigned to OGs, with values ranging from 87.2% in *M. truncatula* to 99.3% in *Lupinus angustifolius*. In *G. max* and *G. soja*, 93.4% and 98.2% of genes, respectively, were grouped into OGs (Figure 1, Figure S1).

**Figure 1:**
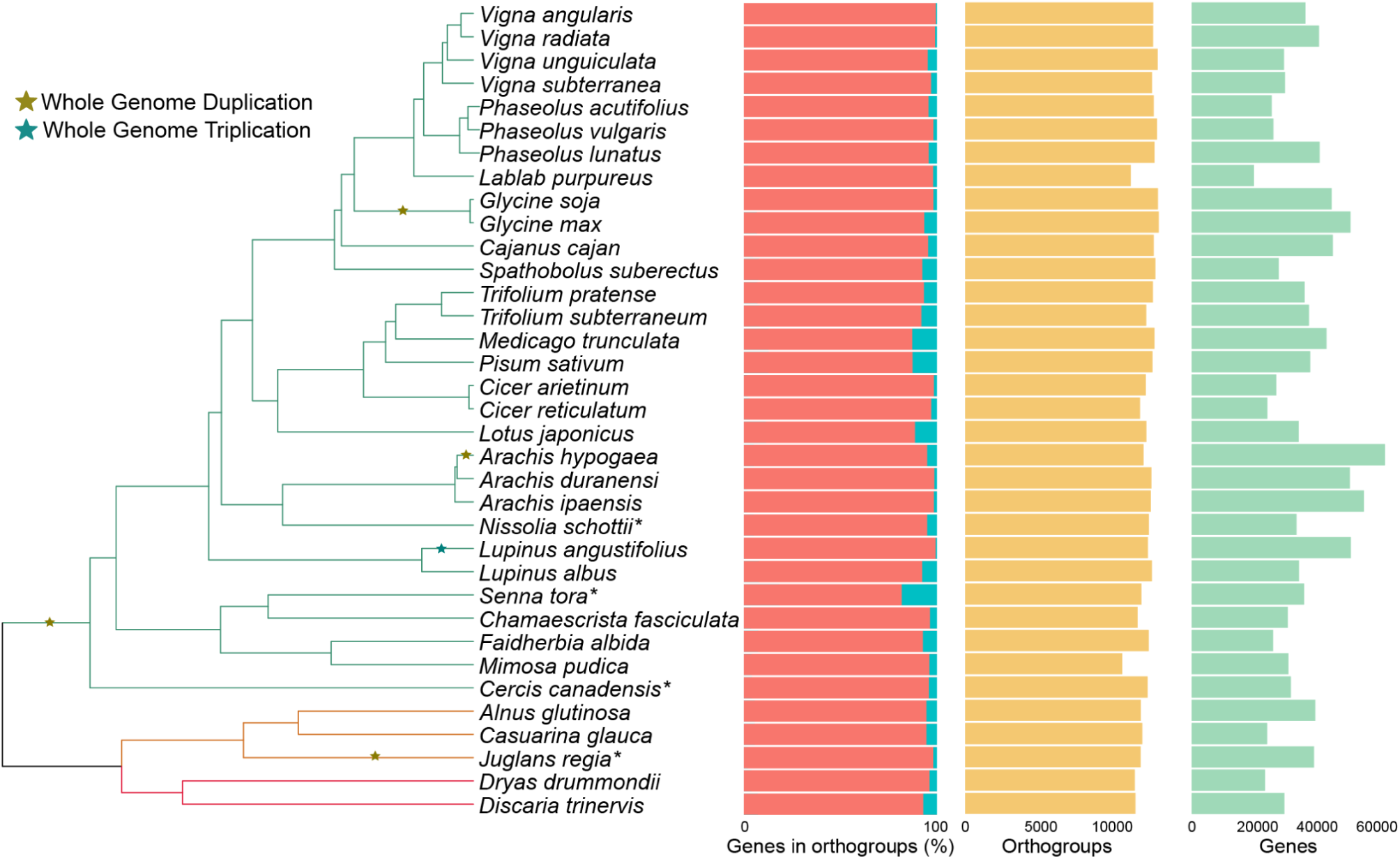
Phylogenetic tree of 35 plant species analyzed in this study. Node colors indicate taxonomic orders: Fabales (green), Fagales (orange), and Rosales (red). The first column displays the percentage of genes assigned to orthologous groups (OGs) per genome, with the non-assigned fraction shown in light blue. The second column represents the number of OGs per genome, and the third column shows the number of non-redundant genes. A detailed version of the tree including inferred gene gains and losses at internal nodes is provided in Figure S1. Branch lengths are not to scale. * Indicates species that lost the ability to form nodules.

Interestingly, approximately 31.5% of the 49,785 OGs were exclusive to a single species. We hypothesize that these species-specific OGs may result from lineage-specific expansions or technical artifacts. To minimize overestimation in gene family expansion/contraction analyses, we restricted our CAFE analyses to a subset of 13,776 OGs that are not species-specific and exhibit suitable variance in gene counts across taxa (see Methods for details). Among the top 50 largest OGs, *A. hypogaea* exhibited the highest average gene copy number (µ = 10.3 members), followed by *G. max* (µ = 10.08 members), *G. soja* (µ = 9.05 members), and *L. angustifolius* (µ = 8.96) (Table S2). These findings are consistent with the occurrence of recent WGDs in these lineages (Flagel & Wendel, 2009; Shoemaker et al., 2006; L. Wang et al., 2019), further supporting the robustness of our approach. Using the time-calibrated phylogenetic tree and the corresponding matrix comprising gene counts for each species across 13,776 OGs, we assessed gene family expansions and contractions. Our analysis revealed a predominance of expansions over contractions throughout legume evolution. Among the 100 largest expanded OGs, *Arachis, Glycine*, and *M. truncatula* showed the most substantial increases in gene family size. Notably, only one of the top 100 most expanded OGs, OG0000135, encoding Aspartic Protease CDR1-related proteins, was functionally linked to nodulation (Figure 2). This result aligns with previous findings in *L. japonicus* (Yamaya-Ito et al., 2018) and *P. vulgaris* (Olivares et al., 2011), which suggest that aspartic proteases play a role in recruiting nitrogen-fixing bacteria from the environment.

**Figure 2:**
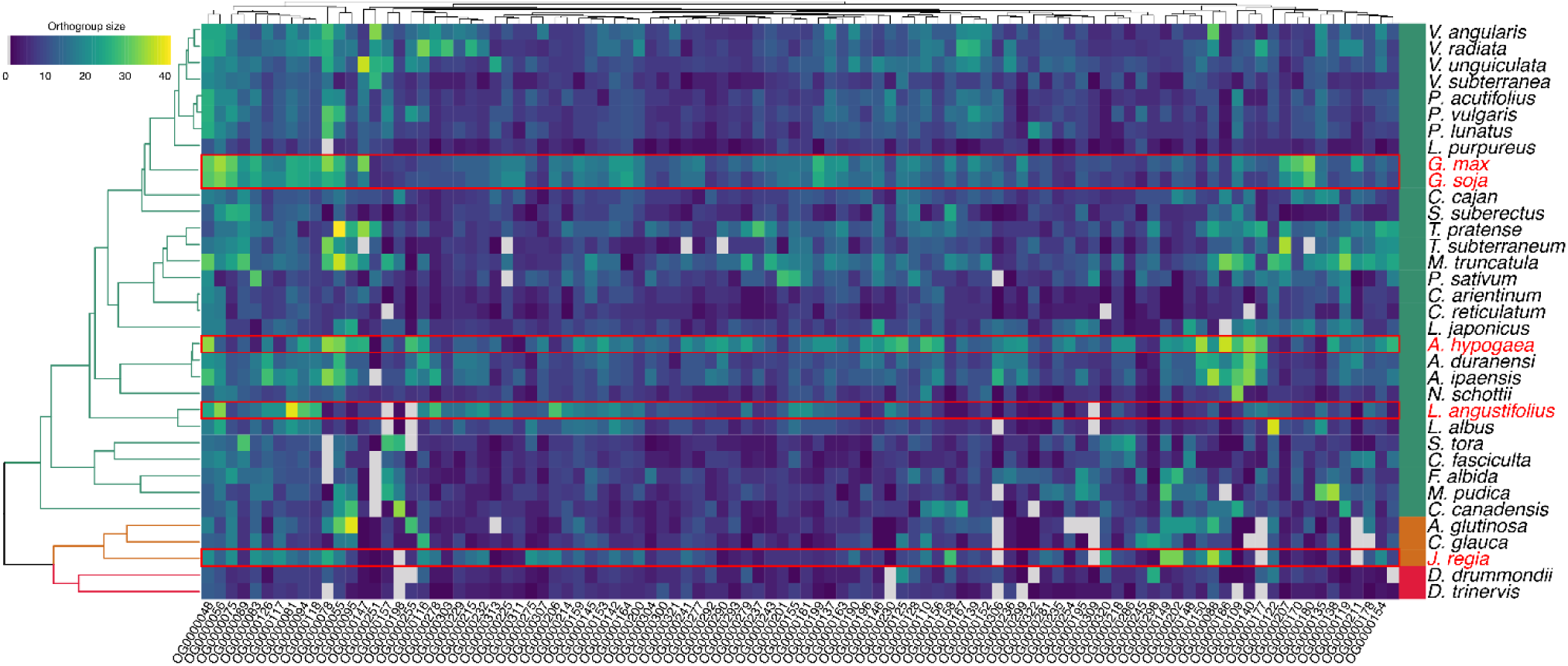
Number of Orthologs in the top 100 expanded gene families. Node colors indicate the order: Fabales (green), Fagales (orange), and Rosales (red). Cell colors represent the number of members of each species within the respective OG; light gray indicates the absence of members in a given species. Species with lineage-specific WGDs in addition to the ancestral WGD shared by Fabaceae are highlighted in red.

The high number of expansions in *G. max* and *G. soja* (Figure 2) could be perceived as a product of the WGD event in the *Glycine* ancestor (Figure 1). Nonetheless, our results reveal a greater gene family expansion in *G. max* compared to *G. soja* (Figure 3), consistent with previous research, which highlight the nuances between cultivated and wild soybean. For instance, Jiang et al. (2022) reported 12,193 expanded families of *G. max* after analyzing 14 plant species, while Varshney et al. (2017) found 6,993 expanded gene families in *G. max* while examining 11 plant species, both figures differing from our total. These discrepancies are likely due to variations in genome assemblies and gene prediction methodologies, which complicate direct cross-study comparisons. Moreover, considering that *G. max* and *G. soja* share the the *Glycine* WGD, we hypothesize that the expansion pattern observed in *G. max* is driven by duplication events other than WGD (in the case of gene gains in *G. max*) and subsequent gene losses in *G. soja* (Figure 3).

**Figure 3.**
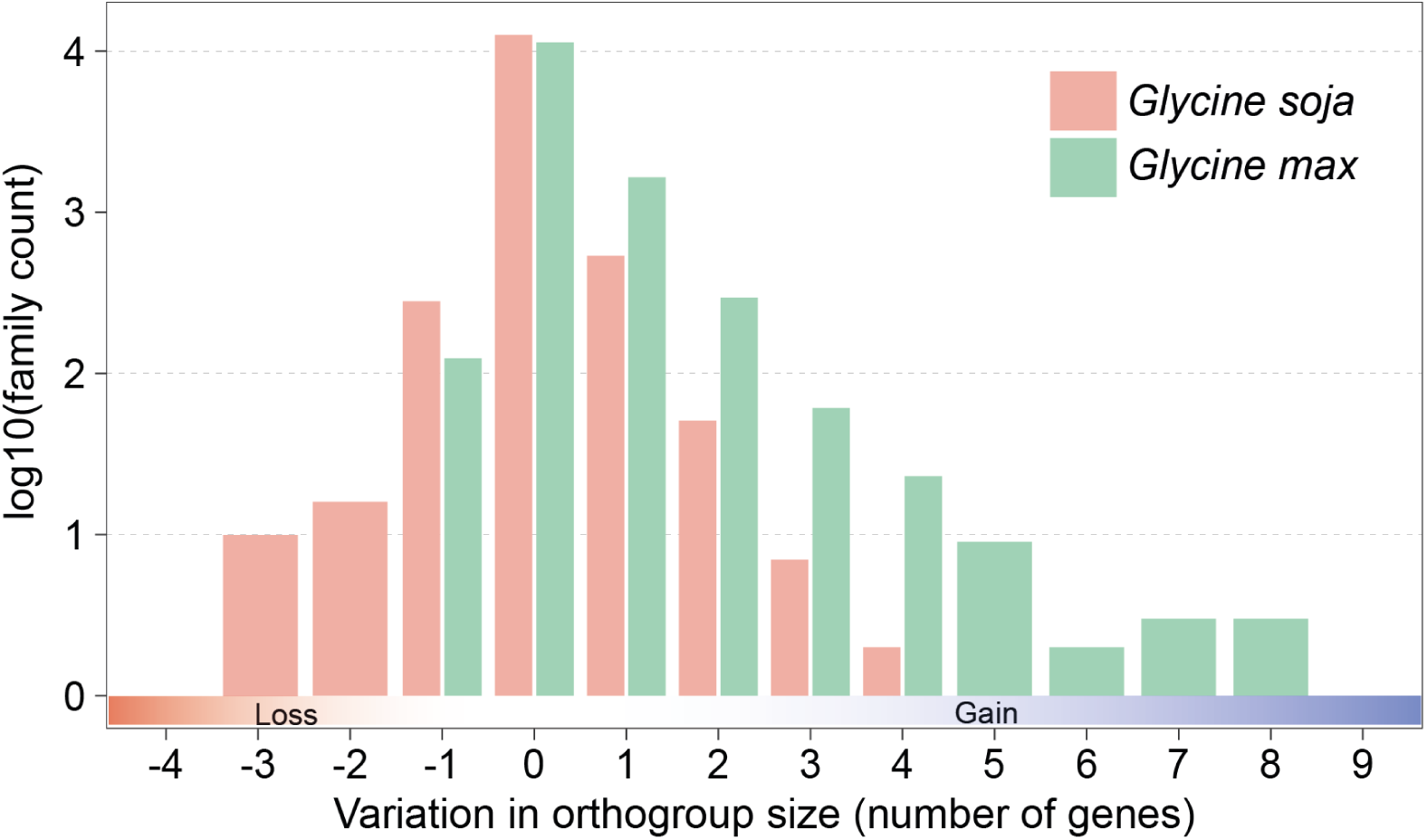
Orthogroup (OG) member changes in *G. max* vs *G. soja*.

Numerous studies have highlighted the pivotal role of WGDs in the expansion of gene families in plants (Hofberger et al., 2015; Hong et al., 2021; Jourda et al., 2014; Lee et al., 2020; Lei et al., 2012; Moharana & Venancio, 2020; Ren et al., 2018). In our analysis, the last common ancestor of the *Glycine* genus exhibited the highest number of expanded families, followed by *A. hypogaea* and *L. angustifolius* (Figure 1). These findings support the hypothesis that the WGD event within the *Glycine* lineage is a major driver of the extensive gene family expansions in soybean (Figure 1, Figure 2), as proposed in other studies (Deshmukh & Datta, 2023; Mengarelli & Zanor, 2021; Sangi et al., 2021; L. Wang et al., 2021, 2021). Given that nodulation predates the divergence of the *Glycine* genus, the genetic framework for nodulation was likely already established in the last common ancestor of Papilionoideae (Mathesius, 2022). However, the extent to which genus- or species-specific recruitment of additional nodulation genes has occurred, and the role of WGD and gene family expansions in fostering innovations within these genetic systems, remains underexplored. In the following sections, we address these questions by integrating a comprehensive RNA-Seq dataset with our gene family analyses.

### Tissue-specific expression pattern of relevant nodulation-related genes in soybean

Nodulation is a complex process involving diverse mechanisms such as receptor-mediated signal transduction, hormonal regulation, nodule organogenesis, autoregulation of nodulation, plant defense, nodule senescence, symbiotic metabolism, and ion transport. In a systematic review, Roy et al. (2020) identified at least 196 genes experimentally validated to participate in these processes in *L. japonicus*, *M. truncatula*, *G. max,* or *P. vulgaris* through forward- and reverse-genetic approaches such as RNAi, CRISPR-Cas9, and transposon-mediated knockouts. We used this comprehensive catalog of nodulation-related genes as a reference to investigate the transcriptional patterns in soybean, aiming to identify candidate genes for further functional studies and crop improvement. To examine gene family dynamics across legumes, we integrated the OGs defined in our study with gene family data from the PLAZA Dicots 5.0 database (Van Bel et al., 2022). While our OGs enabled robust predictions of gene family expansion and contraction across 35 species, cross-referencing with PLAZA, despite including only 10 legume species, provided valuable curated family annotations that enriched our downstream analyses. Through this integrated framework, we found that the 196 nodule-related genes reported by Roy *et al*. were represented by 185 genes in the soybean genome (Table S3). These genes were distributed in 40 OGs and 20 PLAZA families (database accessed on Feb. 2024). Notably, six nodulation-related genes, including two Defective in Nitrogen Fixation (DNF) genes, two Nitrogen Fixation Specificity (NFS) genes, one Small Nodulin Acidic RNA-Binding Protein 2 (SNARP2) gene, and one Interacting Protein DMI3 (IPD3) gene, were assigned to OGs and PLAZA families that lacked *G. max* orthologs, suggesting potential lineage-specific losses in soybean.

The OGs containing soybean members exhibited varied expression profiles across tissues, with a distinct cluster of genes showing strong preferential expression in nodules (Figure 4, Figure S2). Among the 185 nodulation-related genes analyzed, 131 showed expression levels above 1 TPM in nodules, 99 exceeded 5 TPM, and 68 surpassed 10 TPM, based on geometric means of TPM across samples (Table S3). Out of the 20 nodule-related PLAZA gene families, 16 contained at least one member with average nodule expression ≥10 TPM. Notably, 10 of these families contain nodule-specific genes: ORTHO05D000267, ORTHO05D000434, ORTHO05D000845, ORTHO05D001143, ORTHO05D001559, ORTHO05D008022, ORTHO05D016325, ORTHO05D017982, ORTHO05D029738, and ORTHO05D215446. Based on the Soybean Expression Atlas data (Figure S2), 28 genes were classified as nodule-specific, 12 as root- and nodule-specific, and 7 as root-specific. The nodule-specific cluster is enriched in key gene families, including four main families: NLP family proteins, Aspartic Proteinase CDR1, Nodulins, and GRAS/SCARECROW TFs. The nodule-specific cluster encompasses OG0000135 (Figure 2), which is associated with ORTHO05D000267, a PLAZA family that contains three additional OGs (Table S3). OG0000135 ranked among the top 100 expanded OGs in legumes (p-value ≤ 0.05, ranked by expansion magnitude) and includes 15 soybean aspartic-type endopeptidases, such as the nodule-specific genes *Glyma.15G260700*, *Glyma.15G260600*, and *Glyma.08G166200* (Figure 4, Figure S2). Interestingly, several genes in OG0000135 (including the three aforementioned genes) also encode xylanase inhibitor domains (PF14541 and PF14543), which traditionally function in plant defense against herbivores and pathogens (Misas-Villamil & van der Hoorn, 2008). Their expression in nodules raises intriguing questions about their potential roles in modulating the interface between defense and symbiosis. Given the complex interactions in the rhizosphere, xylanase inhibitors may help regulate cell wall remodeling during nodulation while maintaining protective barriers. Other studies suggest that activation of defense genes, including xylanase inhibitors, may be essential for successful nodulation, allowing plants to accommodate beneficial rhizobia while deterring pathogenic microbes (J. Yang et al., 2022). The prominent expansion and nodule-specific expression of aspartic proteases in OG0000135 highlight their potential importance in nodulation and warrant further investigation into their dual roles in symbiotic and defense-related pathways.

**Figure 4:**
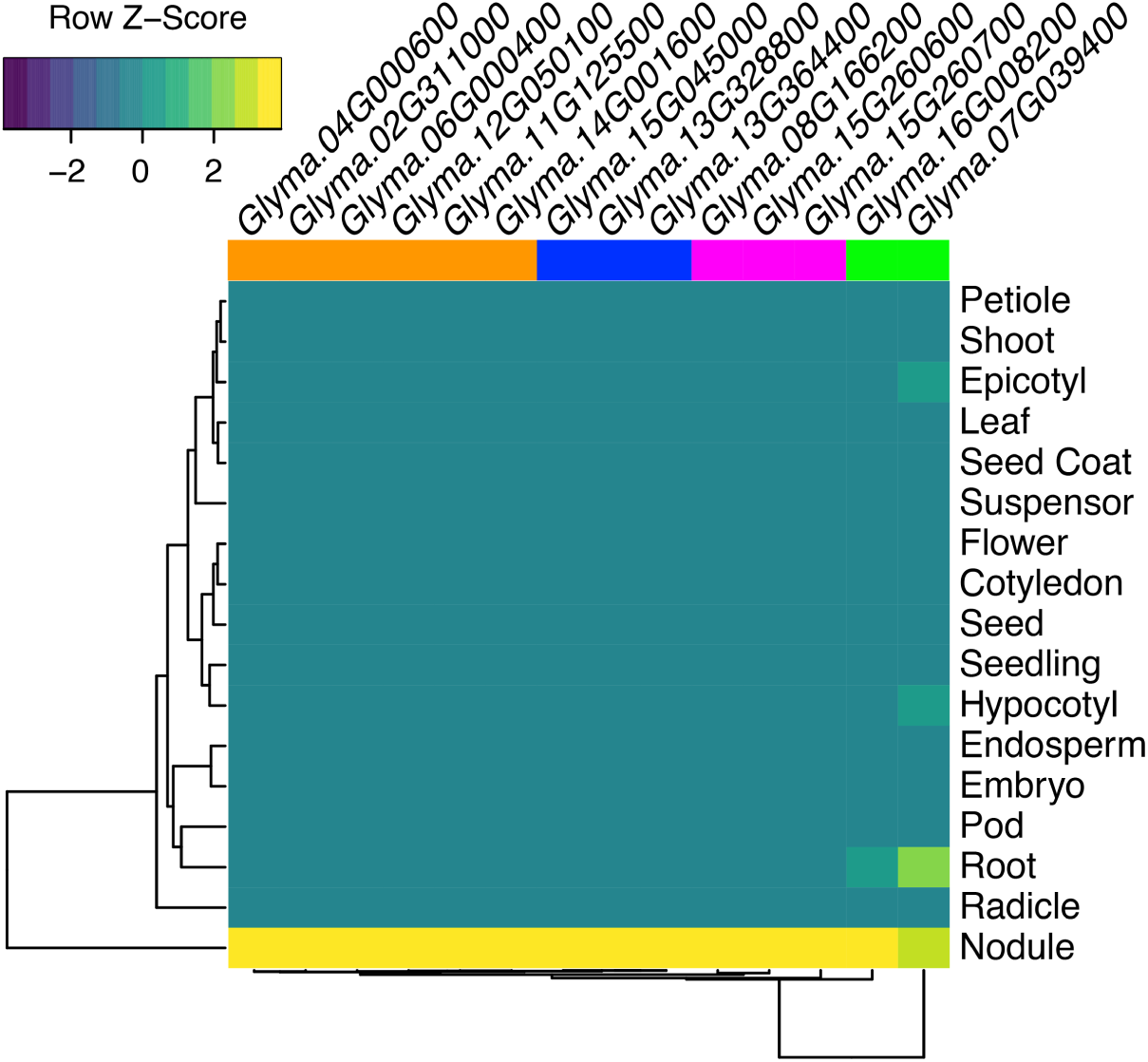
Expression profiles of 14 genes with preferential expression in soybean nodules. This gene set encompasses members of the NLP family (orange), aspartic proteinase CDR1 (magenta), nodulins (blue), and GRAS/SCARECROW transcription factors (green).

Our analysis identified five nodule-specific GRAS/SCARECROW TFs in soybean, distributed across two PLAZA families. Specifically, *Glyma.07G039400* and *Glyma.16G008200* (Figure 4) belong to ORTHO05D008022, while *Glyma.04G251900*, *Glyma.06G110800*, and *Glyma.13G081700* are part of ORTHO05D006939 (Figure S2, Table S3). GRAS TFs play essential roles in plant development, including root and shoot growth as well as the regulation of the shoot apical meristem (Bolle, 2004). The preferential expression of these genes in soybean nodules suggests their functional recruitment for symbiotic processes, highlighting their potential involvement in modulating root architecture and function under both symbiotic and non-symbiotic conditions. This finding aligns with previous studies in *M. truncatula*, which have implicated GRAS TFs in nodulation (Hirsch et al., 2009).

We also identified six nodule-specific members of the NLP family (NLP1-related or NLP8-related, PLAZA Family ORTHO05D000434): *Glyma.12G050100*, *Glyma.04G000600*, *Glyma.02G311000*, *Glyma.06G000400*, *Glyma.14G001600*, and *Glyma.11G125500* (Figure 4, Figure S2). These proteins are defined by the presence of two key domains: the PB1 domain, which facilitates protein-protein interactions such as those between NLP1 and NIN in *M. truncatula* to modulate nodule development under varying nitrate conditions (Lin et al., 2018), and the RWP-RK domain, which is crucial for regulating cellular responses to nitrogen availability (Chardin et al., 2014). In nodulating species, RWP-RK acts as a TF that activates genes essential for nodulation by mediating symbiont recognition of symbiotic and infection thread formation (Amin et al., 2023). This dual functionality highlights the role of NLP proteins in integrating environmental nitrogen sensing with developmental processes critical for nodulation. As expected, nodulins exhibited exclusive and high expression levels in nodules, with prominent examples such as *Glyma.15G045000*, *Glyma.13G364400*, and *Glyma.13G328800* (Figure 4, Figure S2). Nodulins have long been recognized as central components in the establishment and maintenance of symbiont colonization (Sanchez et al., 1991). While most nodulation genes display strong expression in nodules, many also show activity in other tissues, reflecting the systemic nature of nodulation signaling and its integration with broader plant developmental and physiological processes.

### Expansion and recruitment of β-glucosidase genes to nodulation in soybean

We also identified novel candidate genes putatively involved in nodulation by integrating expanded families identified by CAFE with the large gene expression dataset from the Soybean Expression Atlas and comparison with the *M. truncatula* expression profiles. This analysis revealed 262 gene families significantly expanded in the *Glycine* genus (p-value ≤ 0.05), comprising a total of 2,225 genes (Table S4). Of these, 49 genes from 38 families showed at least 10-fold higher expression in nodules compared to other tissues (Table 1). Notably, three PLAZA families, ORTHO05D000033, ORTHO05D000040, and ORTHO05D000267, stood out for harboring the largest number of nodule-preferentially expressed members, each with three genes.

**Table 1:** Expanded gene families with their respective members with higher expression in nodules. GM: geometric mean.

| Soybean Gene | Plaza Family | TPM (nodule) | TPM (GM) | Fold-Change | Function |
| --- | --- | --- | --- | --- | --- |
| *Glyma.01G078300* | ORTHO05D000040 | 512.564 | 1.027 | 498.90 | Cytochrome P450 71B21-related |
| *Glyma.08G076800* | ORTHO05D001343 | 440.462 | 1.139 | 386.80 | EamA-like Transporter Family (EamA) |
| *Glyma.13G191600* | ORTHO05D000117 | 343.264 | 1.033 | 332.21 | Sulfotransferase Sult |
| *Glyma.18G247100* | ORTHO05D000108 | 229.726 | 1.031 | 222.87 | Glucosyl/Glucuronosyl Transferases |
| *Glyma.13G247000* | ORTHO05D009876 | 446.458 | 2.502 | 178.47 | Wound-induced Protein (DUF3774) |
| *Glyma.14G064400* | ORTHO05D000270 | 173.136 | 1.059 | 163.43 | Proprotein Convertase Subtilisin/Kexin |
| *Glyma.18G263700* | ORTHO05D000159 | 163.896 | 1.126 | 145.54 | Licodione 2'-O-methyltransferase |
| *Glyma.11G222300* | ORTHO05D000312 | 821.875 | 6.291 | 130.65 | Aspartic Proteinase CDR1-Related |
| *Glyma.15G260700* | ORTHO05D000267 | 105.188 | 1.027 | 102.44 | Aspartic Proteinase CDR1-Related |
| *Glyma.06G286600* | ORTHO05D000154 | 186.395 | 2.123 | 87.78 | O-methyltransferase |
| *Glyma.01G195700* | ORTHO05D001581 | 150.484 | 1.817 | 82.80 | Growth Factor Protein-related |
| *Glyma.17G125400* | ORTHO05D000040 | 111.980 | 1.465 | 76.43 | Costunolide Synthase |
| *Glyma.15G260600* | ORTHO05D000267 | 45.443 | 1.015 | 44.78 | Aspartic Proteinase CDR1-Related |
| *Glyma.13G118800* | ORTHO05D000084 | 75.949 | 1.697 | 44.75 | ABC Transporter B Family Member 11-related |
| *Glyma.08G166200* | ORTHO05D000267 | 39.742 | 1.047 | 37.96 | Aspartic Proteinase CDR1-related |
| *Glyma.12G191700* | ORTHO05D001793 | 91.472 | 2.465 | 37.10 | Cellulose Synthase-Like Protein B1-Related |
| *Glyma.19G251500* | ORTHO05D000734 | 38.885 | 1.171 | 33.19 | Cucumisin |
| *Glyma.02G018700* | ORTHO05D000033 | 102.299 | 3.215 | 31.82 | Hydroxyisourate Hydrolase / HIUHase |
| *Glyma.05G126900* | ORTHO05D001454 | 102.125 | 3.302 | 30.93 | Stress Up-regulated Nod 19 (SURNod19) |
| *Glyma.15G251800* | ORTHO05D000264 | 52.471 | 1.732 | 30.29 | Glutathione S-Transferase |
| *Glyma.03G104500* | ORTHO05D000052 | 39.883 | 1.336 | 29.85 | Hydroquinone:O-glucosyltransferase |
| *Glyma.10G051700* | ORTHO05D003343 | 26.120 | 1.201 | 21.75 | Leucine-rich Repeat-containing Protein |
| *Glyma.17G125300* | ORTHO05D000040 | 26.260 | 1.264 | 20.78 | Cytochrome P450 CYP2 Subfamily |
| *Glyma.08G071200* | ORTHO05D000406 | 24.006 | 1.157 | 20.75 | Taurine-transporting ATPase |
| *Glyma.09G014400* | ORTHO05D000141 | 110.446 | 5.330 | 20.72 | Lipid-Transfer Protein/<br>Seed Storage 2S Albumin Superfamily Protein |
| *Glyma.13G221900* | ORTHO05D000255 | 33.671 | 1.640 | 20.53 | Serine Carboxypeptidase-like Clade I (SCPL-I) |
| *Glyma.18G263200* | ORTHO05D000102 | 20.158 | 1.041 | 19.36 | ABC Transporter C Family Member 3-Related |
| *Glyma.07G136100* | ORTHO05D000312 | 28.410 | 1.535 | 18.51 | Tropinone Reductase I |
| *Glyma.15G049800* | ORTHO05D001343 | 42.568 | 2.461 | 17.30 | EamA-like Transporter Family (EamA) |
| *Glyma.13G252600* | ORTHO05D000121 | 28.815 | 1.676 | 17.20 | Cysteine-rich Secretory Protein-related |
| *Glyma.09G260500* | ORTHO05D012095 | 16.679 | 1.017 | 16.41 | Bowman-Birk Serine Protease Inhibitor Family |
| *Glyma.17G052900* | ORTHO05D000051 | 57.347 | 3.717 | 15.43 | Peroxidase 59 |
| *Glyma.06G315400* | ORTHO05D000256 | 23.807 | 1.561 | 15.25 | Protein of Unknown Function (DUF640) |
| *Glyma.07G151900* | ORTHO05D000033 | 24.681 | 1.657 | 14.89 | Glycosyl Hydrolase/<br>β-glucosidase 1-related |
| *Glyma.07G089700* | ORTHO05D000450 | 16.191 | 1.092 | 14.83 | Abieta-7,13-dien-18-ol Hydroxylase /<br>CYP720B1 |
| *Glyma.11G185200* | ORTHO05D000338 | 26.553 | 1.869 | 14.21 | Long-chain-fatty-acyl-CoA Reductase/<br>Alcohol-forming Fatty Acyl-CoA Reductase |
| *Glyma.08G021100* | ORTHO05D001215 | 685.130 | 48.350 | 14.17 | Protein of Unknown Function (DUF1070) |
| *Glyma.10G168300* | ORTHO05D000134 | 27.120 | 1.948 | 13.92 | Protein of Unknown Function (DUF239) |
| *Glyma.20G003600* | ORTHO05D000159 | 13.485 | 1.007 | 13.39 | Licodione 2'-O-methyltransferase |
| *Glyma.08G189500* | ORTHO05D000130 | 48.443 | 3.656 | 13.25 | Linoleate 9S-lipoxygenase (LOX1_5) |
| *Glyma.18G274400* | ORTHO05D000083 | 30.818 | 2.337 | 13.19 | UDP-glucosyl Transferase 73C |
| *Glyma.15G119100* | ORTHO05D000141 | 67.291 | 5.145 | 13.08 | Lipid-Transfer Protein/<br>Seed Storage 2S Albumin Superfamily Protein |
| *Glyma.13G066100* | ORTHO05D000085 | 42.701 | 3.426 | 12.46 | AAA-type ATPase Family Protein-related |
| *Glyma.10G159700* | ORTHO05D000468 | 46.631 | 3.759 | 12.40 | Bifunctional L-3-cyanoalanine Synthase/<br>Cysteine Synthase D1-related |
| *Glyma.17G053000* | ORTHO05D000051 | 36.381 | 2.960 | 12.29 | Peroxidase 59 |
| *Glyma.18G005000* | ORTHO05D000526 | 104.133 | 8.984 | 11.59 | Protein of Unknown Function (DUF3339) |
| *Glyma.16G097200* | ORTHO05D000033 | 24.304 | 2.166 | 11.22 | Glycosyl Hydrolase/<br>β-glucosidase 1-related |
| *Glyma.16G017200* | ORTHO05D000932 | 110.695 | 10.468 | 10.57 | Inorganic Pyrophosphatase F11O4.12 |
| *Glyma.08G095300* | ORTHO05D000166 | 10.401 | 1.027 | 10.13 | Protein Agamous-like 82 |

To better understand the potential recruitment of the ORTHO05D000033 gene family of β-glucosidases for nodule-related functions, we examined the expression of the *M. truncatula* orthologs therein. This analysis was guided by the hypothesis that certain genes may have been recruited for nodulation before the divergence of *M. truncatula* and soybean, estimated at approximately 45 MYA (adjusted time, TimeTree Database) (Kumar et al., 2017). Using a SSN, we identified three distinct clusters and seven unclustered genes within the ORTHO05D000033 family (Figure 5). A similarity cutoff of 47% identity was applied, enabling the formation of a subcluster that revealed distinct expression patterns between *G. max* and *M. truncatula*. Interestingly, one of these clusters revealed contrasting expression profiles between the two species: *M. truncatula* orthologs did not show nodule-preferential expression, whereas four soybean genes, *Glyma.07G151900*, *Glyma.16G097200*, *Glyma.02G018700*, and *Glyma.02G155800*, exhibited high nodule-specific expression (z-scores ∼2). As proposed by Gerlt (2017), SSN clustering can serve as a proxy for functional similarity, helping to infer recruitment and neofunctionalization events. These findings support the notion that the nodule-specific expression of these soybean genes likely emerged after the divergence from *M. truncatula*, reflecting evolutionary specialization within the ORTHO05D000033 family.

**Figure 5:**
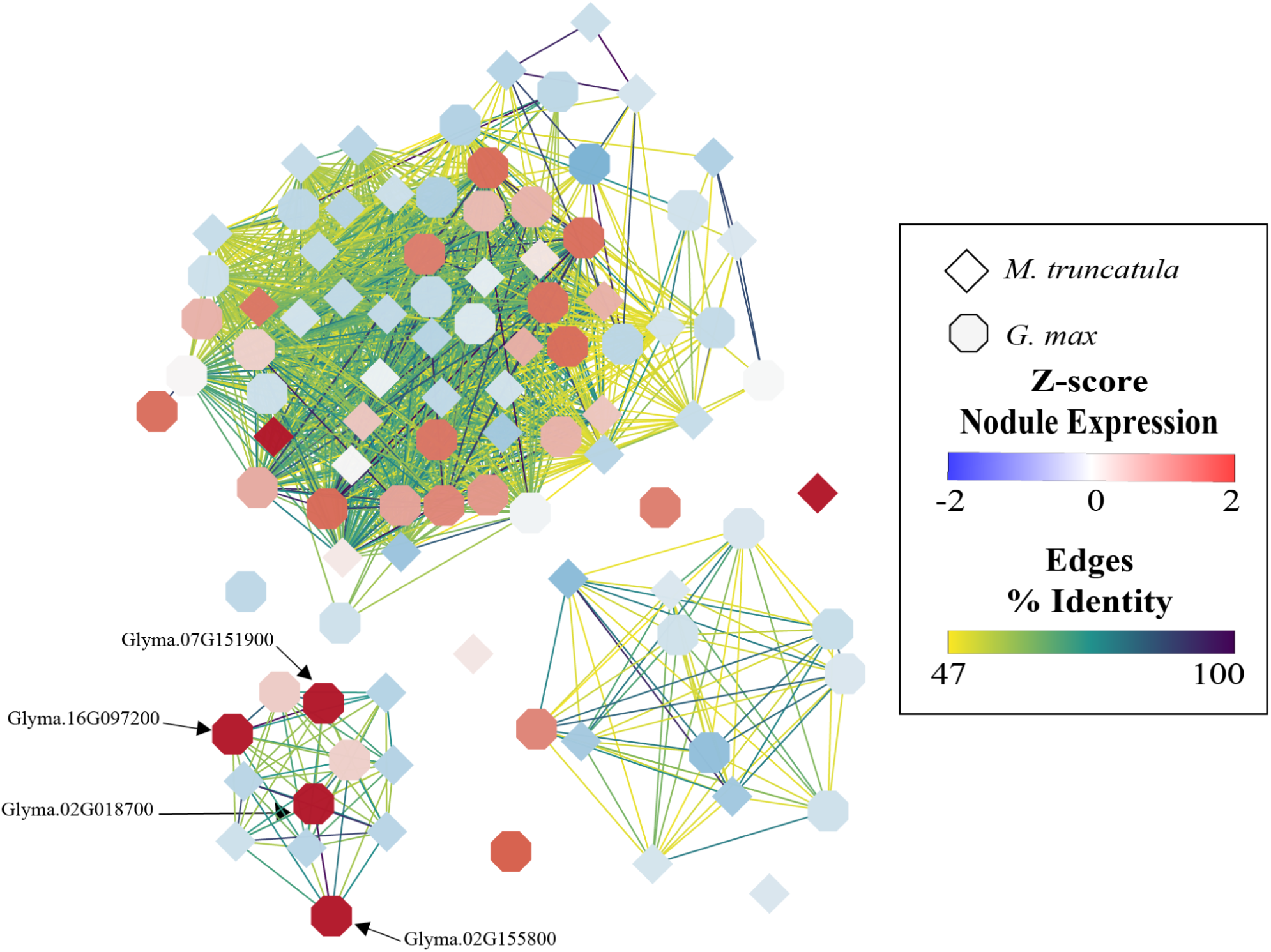
Sequence Similarity Network of a β-glucosidase gene family (ORTHO05D000033), represented by 49 members in soybean (hexagonal nodes) and 47 members in *M. truncatula* (diamond nodes). The figure depicts three putative isofunctional clusters based on the similarity among family members. Four members in soybean (labeled nodes) are part of a new cluster and are preferentially expressed in nodules.

Considering the divergent expression pattern of the *G. max* and *M. truncatula* orthologs in this subcluster, we raised the question of whether these genes are a product of WGD events in legumes, in the Glycine genus or another independent event. To try to describe the origin of these four β-glucosidase genes, we performed a Ka/Ks ratio analysis. Although the results were inconclusive in determining the exact duplication order among the genes, they offered insights into the timing of these duplication events. Specifically, divergence time estimates suggest two possible scenarios for the origins of *Glyma.16G097200* (Table 2). This gene may have arisen either from an ancient segmental duplication involving *Glyma.07G151800* or from a more recent dispersed duplication of *Glyma.07G151900*, which is a neighboring gene of *Glyma.07G151800* (tandem duplication ∼22.13 MYA). The estimated age of the tandem duplication predates the well-characterized WGD in the *Glycine* genus (∼13 MYA), indicating that these duplication events are part of an ongoing evolutionary trajectory independent of the WGD (Table 2). This is, to our knowledge, the first study to report the involvement of this gene family in nodulation. While β-glucosidases are traditionally associated with the hydrolysis of various glycosides and oligosaccharides, they are also known to participate in plant defense (Ma et al., 2024). The preferential and high expression of these genes in nodules, together with their evolutionary recruitment post-WGD, suggests a dual role in both symbiotic interactions and defensive responses. These findings support the emerging view that certain metabolic enzymes may be co-opted into symbiosis-related pathways, balancing microbial accommodation with protective functions in the rhizosphere.

**Table 2:** Ka/Ks analysis and duplication types for *Glyma.07G151900, Glyma.16G097200, Glyma.02G018700,* and *Glyma.02G155800*.

| Duplicated gene pairs | Ka | Ks | Ka/Ks | Time (MYA) | Duplication Type |
| --- | --- | --- | --- | --- | --- |
| *Glyma.16G097200/Glyma.07G151800* | 0.0823012 | 0.262995 | 0.312938 | 21.31 | Segmental |
| *Glyma.07G151900/Glyma.07G151800* | 0.0822666 | 0.271775 | 0.302702 | 22.13 | Tandem |
| *Glyma.02G018700/Glyma.02G155800* | 0.002171 | 0.003012 | 0.720678 | 0.24 | Dispersed |
| *Glyma.14G211700/Glyma.02G155800* | 0.105512 | 0.408681 | 0.258177 | 33.44 | Dispersed |
| *Glyma.02G018700/Glyma.14G211700* | 0.116868 | 0.429562 | 0.272063 | 35.24 | Dispersed |
| *Glyma.07G151900/Glyma.16G097200* | 0.00274348 | 0.014189 | 0.193352 | 0.82 | Dispersed |

The ORTHO05D000040 family comprises three soybean cytochrome P450 genes, *Glyma.01G078300*, *Glyma.17G125300*, and *Glyma.17G125400*, that exhibit exceptionally high expression levels in nodules (Table S3). All three genes were annotated by InterProScan as part of the Cytochrome P450, E-class group I family (IPR002401). Notably, their expression levels in nodules were at least 20-fold higher than in other tissues. For example, *Glyma.01G078300* showed a striking expression of ∼500 TPM in nodules, with negligible expression elsewhere (TPM values < 1). The cytochrome P450 family is one of the largest and most functionally diverse in plants, contributing to the biosynthesis of a broad range of secondary metabolites such as pigments, flavor compounds, and defense-related molecules (Guengerich, 2012). The family has undergone substantial expansion across plant lineages (Nelson et al., 2008), and while their roles in various physiological processes have been investigated (Guttikonda et al., 2010; Z. Yang et al., 2023), this is, to our knowledge, the first study to associate *Glyma.01G078300*, *Glyma.17G125300*, and *Glyma.17G125400* with nodulation. These findings point to a potentially novel role for this cytochrome P450 subfamily in the symbiotic processes underlying nodulation. Their strong and nodule-specific expression suggests a specialized function that may involve modulation of plant metabolism or defense during the establishment and maintenance of rhizobial symbiosis, warranting further functional characterization.

In addition to the genes previously discussed, several others listed in Table 1 are reported here as being associated with nodulation, despite their known involvement in other metabolic pathways. For example, according to Phytozome annotations (available at https://phytozome-next.jgi.doe.gov/), *Glyma.17G052900* and *Glyma.17G053000* are involved in baicalein degradation. *Glyma.03G104500* is linked to arbutin biosynthesis, and *Glyma.18G274400* plays a role in saponin biosynthesis, processes that are typically not linked to nodulation. The expression of these genes in nodules may reflect broader or multifunctional roles, possibly related to secondary metabolism, stress response, or plant-microbe signaling. It remains unclear whether their nodule-specific expression is condition-dependent (e.g., influenced by developmental stage or environmental cues), or whether they contribute to functions beyond organogenesis or nitrogen fixation. These findings underscore the complexity of the nodulation process and call for further functional validation studies to dissect the precise roles of these genes within the nodule microenvironment.

## CONCLUSIONS

In this study, we provide new insights into soybean genes involved in nodulation by integrating gene family expansion analyses with large-scale transcriptomic data. Our results both confirm and extend previous research by corroborating the involvement of aspartic proteases, NLP proteins, nodulins, and GRAS TFs in nodulation. In addition, we report the identification of three cytochrome P450 (E-class group I) and four β-glucosidase genes that are likely associated with nodulation. These genes belong to OGs that have undergone significant expansions in *G. max* and show preferential expression in nodules, highlighting their potential roles in nodulation-related processes.

## CONFLICT OF INTEREST

The authors declare that the research was conducted in the absence of any commercial or financial relationships that could be construed as a potential conflict of interest.

## AUTHOR CONTRIBUTIONS

**Cláudio Benício Cardoso-Silva**: Data curation, Methodology, Formal analysis, Writing – original draft, Writing – review & editing. **Gabriel Quintanilha-Peixoto**: Methodology, Formal analysis, Writing – original draft, Writing – review & editing. **Dayana Kelly Turquetti-Moares:** Formal analysis, Writing – original draft, Writing – review & editing. **Fabricio Almeida-Silva**: Formal analysis, Writing – review & editing. **Thiago M. Venancio**: Conceptualization, Funding acquisition, Project administration, Writing – original draft, Writing – review & editing.

## ACKNOWLEDGMENTS

We acknowledge funding from Fundação Carlos Chagas Filho de Amparo à Pesquisa do Estado do Rio de Janeiro (FAPERJ; grants E-26/201.808/2020, E-26/202.354/2021 and E-26/200.026/2024), Coordenação de Aperfeiçoamento de Pessoal de Nível Superior, Brazil (CAPES; Finance Code 001), and Conselho Nacional de Desenvolvimento Científico e Tecnológico (grants 313024/2023-5, 440221/2022-6, and314347/2020-8). FA-S acknowledges funding from Ghent University (Methusalem funding, BOF.MET.2021.0005.01). The funding agencies had no role in the design of the study and collection, analysis, and interpretation of data, and writing.

## DATA AVAILABILITY

Details regarding the genomes exploited for phylogenomic analysis can be found in Table S1. The RNA-Seq data for *Glycine max* is openly accessible through the Soybean Atlas (available at https://soyatlas.venanciogroup.uenf.br). Additionally, RNA-Seq data for *Medicago truncatula* were sourced from the NCBI SRA repository under Bioproject numbers PRJEB43588 and PRJNA832760.

## Notes

### Competing Interest Statement

The authors have declared no competing interest.

