## Supplementary figures for "Gene family expansions and nodule-specific expression patterns reveal the recruitment of Beta-Glucosidases and Cytochrome P450 genes to nodulation in soybean"

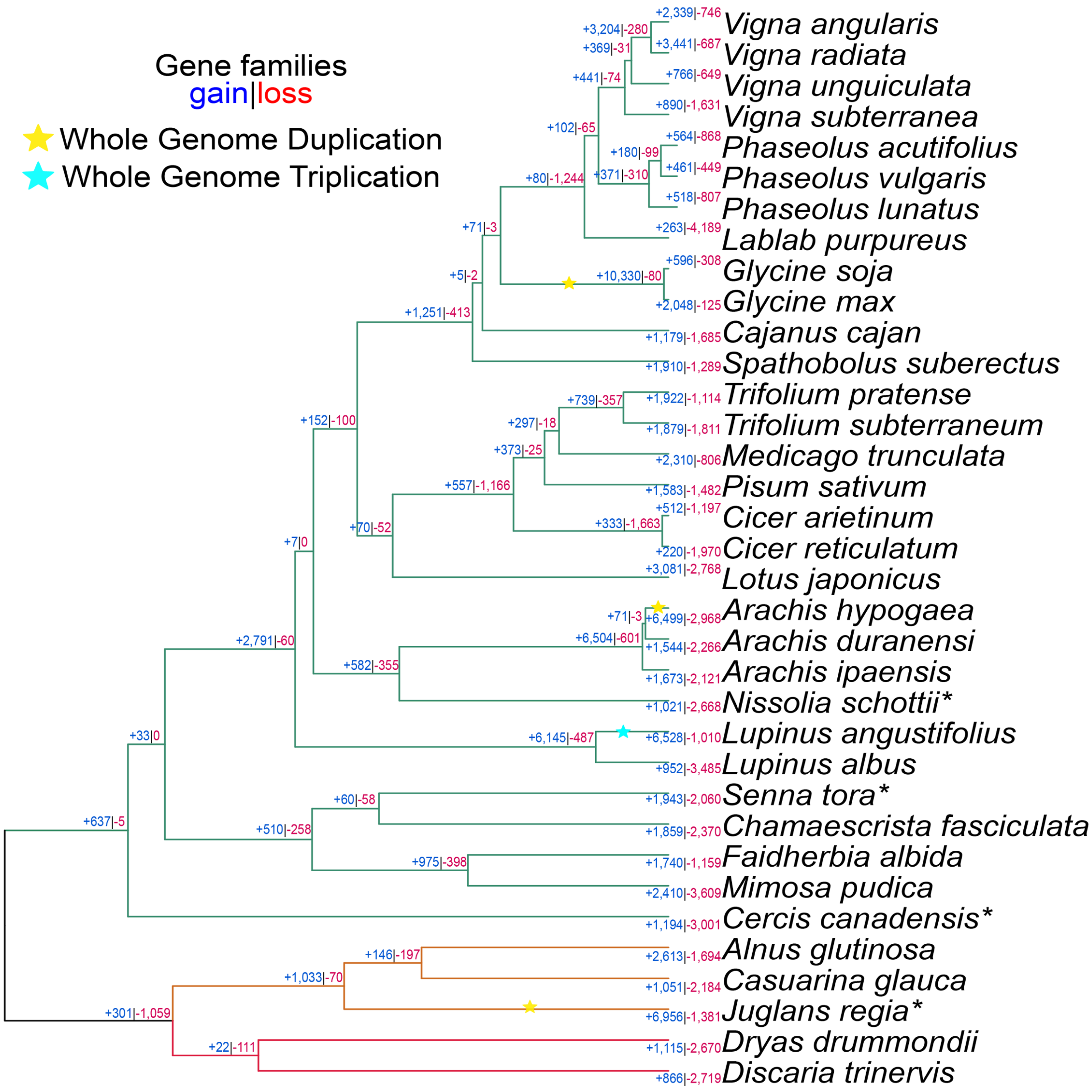


**Figure S1:** Cladogram of the 35 plant species analyzed in this study. Node colors indicate taxonomic orders: Fabales (green), Fagales (orange), and Rosales (red). Gene family contractions and expansions inferred by CAFE 5 are shown in blue and green, respectively. Stars on branches denote whole-genome duplication (WGD) events, while asterisks ("*") next to species names indicate loss of nodulation ability.

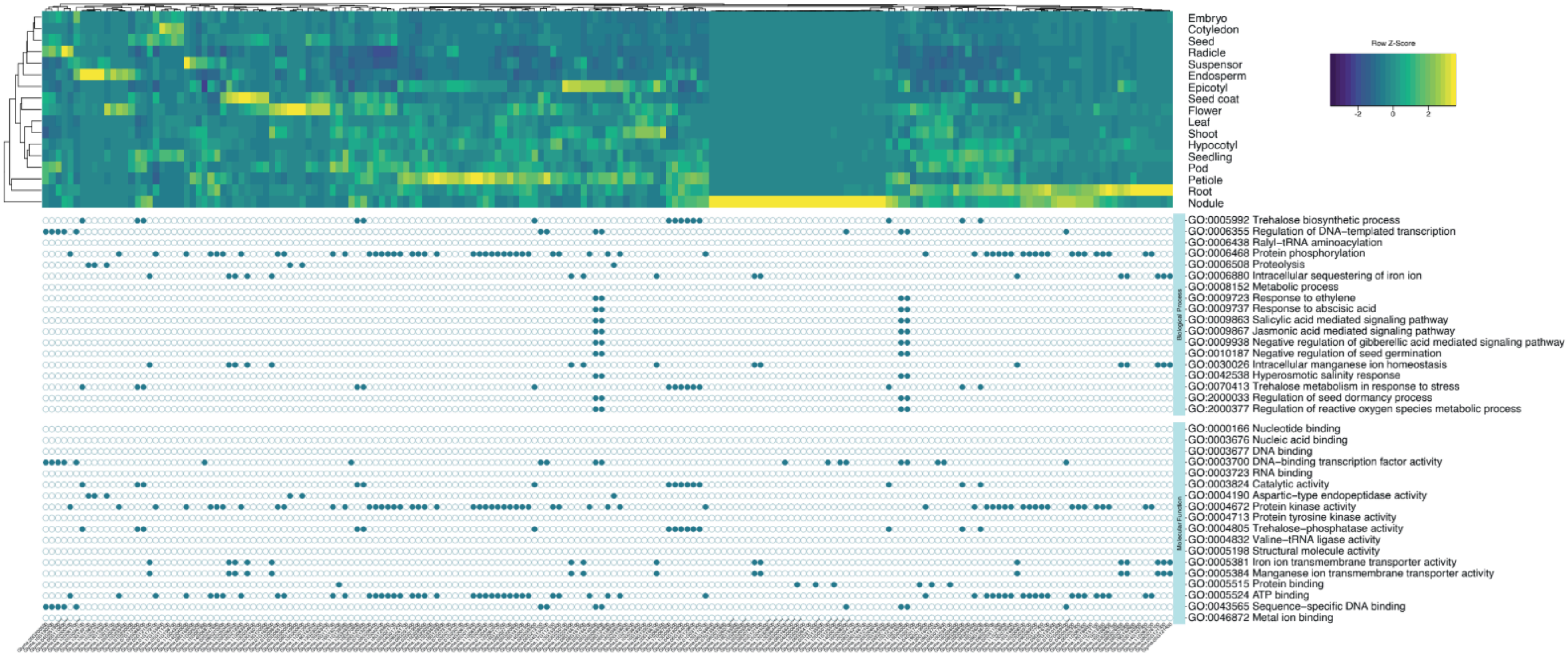


**Figure S2:** Detailed expression and GO annotation of soybean genes associated with nodulation.
